# Improved protein interaction models predict differences in complexes between human cell lines

**DOI:** 10.1101/2024.10.25.620244

**Authors:** Gary R. Wilkins, Jose Lugo-Martinez, Robert F. Murphy

## Abstract

The interactions of proteins to form complexes play a crucial role in cell function. Data on protein-protein or pairwise interactions (PPI) typically come from a combination of sample separation and mass spectrometry. Since 2010, several extensive, high-throughput mass spectrometry-based experimental studies have dramatically expanded public repositories for PPI data and, by extension, our knowledge of protein complexes. Unfortunately, challenges of limited overlap between experiments, modality-oriented biases, and prohibitive costs of experimental reproducibility continue to limit coverage of the human protein assembly map, both underscoring the need for and spurring the development of relevant computational approaches.

Here, we present a new method for predicting the strength of protein interactions. It addresses two important issues that have limited past PPI prediction approaches: incomplete feature sets and incomplete proteome coverage. For a given collection of protein pairs, we fused data from heterogeneous sources into a feature matrix and identified the minimal set of feature partitions for which a non-empty set of protein pairs had complete values. For each such feature partition, we trained a classifier to predict PPI probabilities. We then calculated an overall prediction for a given protein pair by weighting the probabilities from all models that applied to that pair. Our approach accurately identified known and highly probable PPI, far exceeding the performance of current approaches and providing more complete proteome coverage. We then used the predicted probabilities to assemble complexes using previously-described graph-based tools and clustering algorithms and again obtained improved results. Lastly, we used features for three human cell lines to predict PPI and complex scores and identified complexes predicted to differ between those cell lines.

## 1. INTRODUCTION

The interactions of proteins play a crucial role in cellular function. Determining the protein-protein interactions (PPI) underlying all cellular physiology is essential to understanding biological systems at the molecular level. Protein interactions may be direct, in which a pair of proteins physically bind together, or indirect, through being bound to other members of a complex. Support for direct interactions is demonstrated through well-established experimental assays such as yeast two-hybrid (Fields & Song, 1989), whereas indirect interactions are inferred from identifying proteins present in a complex together. For example, in affinity-purification, molecular tags are attached to proteins of interest and after extraction any associated proteins are identified. Alternatively, samples are separated by their physical properties and the frequencies of co-fractionating proteins measured (Dunkley et al., 2004; Foster et al., 2006). In both methods, mass-spectrometry is most frequently used for protein identification.

Advances in high-throughput technology have enabled extending coverage of the human interactome, the set of all PPI, to greater than 1.3 million interactions (Ewing et al., 2007; Hein et al., 2015; Huttlin et al., 2015; Malovannaya et al., 2011; Petrey et al., 2023; Wan et al., 2015). Nonetheless, this is a very small fraction of all PPI expected. CORUM, the dominant resource and reference for mammalian protein complexes, now contains 26% of the human proteome and reports 5,204 human complexes out of an estimated 650,000 complexes (Giurgiu et al., 2019; Stumpf et al., 2008; Tsitsiridis et al., 2023). Moreover, there are significant limitations to the physical processes governing existing experimental methods. Affinity purification requires interactions strong enough to persist through fractionation, and is therefore biased toward false negatives, resulting in missed interactions. Inherent to its design, results are limited to interactions with the tagged protein used for purification (Elhabashy et al., 2022; Low et al., 2021). Results for co-fractionation are limited by insufficient resolving power between organelles or complexes with similar physical properties, and potentially by heterogeneity in those properties among organelles. This adds uncertainty to the identification and subsequent accuracy assessments of predicted PPI and complexes (Skinnider et al., 2023).

In confronting these challenges, recent strategies have been developed to integrate heterogeneous experimental molecular data to predict protein-protein interactions (PPI) and resulting complexes with greater confidence and accuracy (Adamcsek et al., 2006; Altaf-Ul-Amin et al., 2006; Bader & Hogue, 2003; Elhabashy et al., 2022; Liu et al., 2008; Ma et al., 2017; Nepusz et al., 2012; Ou-Yang et al., 2017; Pan et al., 2023; Peng et al., 2014; Qi et al., 2008; Skinnider et al., 2023; Wu et al., 2009). These approaches typically involve calculating a confidence score for each pair or potential interaction, clustering pairs into complexes based on these scores, and evaluating predicted complexes against known and suspected complexes. For example, Drew et al., 2017 trained a predictive model for protein interaction probabilities using three existing data sources and developed a model of interactions between bait-sharing prey, thereby enabling the prediction of novel interactions (Hein et al., 2015; Huttlin et al., 2015; Wan et al., 2015). Lugo-Martinez et al., 2019 explored the hypothesis that experimental measurements that do not directly reflect protein interactions could enhance PPI predictions by incorporating negative evidence, such as lack of co-expression or dissimilar subcellular localization. Similar addition of co-expression and co-localization features resulted in improved predictive accuracy and broader proteome coverage (Elhabashy et al., 2022). Drew et al., 2021 updated their prior protein complex map by integrating over 15,000 proteomic experiments, representing the broadest coverage of human proteins at the time. Additionally, Skinnider et al., 2023 emphasized the importance of co-fractionation mass spectrometry for mapping protein states and interactions, leveraging machine learning for the analysis of co-fractionation profiles to improve the accuracy of interaction predictions.

When constructing models by integrating data from different sources, a major issue can be how to handle feature values that are missing for particular samples. This issue is particularly relevant for predicting PPI since complete sets of features are available for only a very small fraction of protein pairs, making statistical machine learning approaches such as feature inference particularly difficult. We propose an ensemble approach in which different predictors are trained for different feature partitions using samples for which all of the features in that partition is available. We demonstrate that this provides improved predictive accuracy compared to previous approaches and that incorporating additional data sources improves accuracy even further. Lastly, we use features for particular cell lines to make specific predictions for each line and identify differences in predicted PPI and protein complexes between lines.

## 2. METHODS

### 2.1 Predictive model design

For this study, we developed an approach, depicted in Figure 1, to handle the extreme variation in features available for different protein pairs. We refer to this method as Ensemble Learning for Complete Feature Subsets (ELCFS). It begins by identifying groups of observations (protein pairs) for which a subset of the features did not have any missing values. This, in turn, enables the creation of feature partitions that would allow prediction for all pairs without any feature imputation (Figure 1 ii-b). Random forest models (from Scikit-Learn (Pedregosa et al., 2011)) are then learned, using a training set, for each partition. Predictions for the training set are used to estimate the accuracy of the model for each partition; and predicted scores for each test pair are calculated as the accuracy-weighted average of the predictions of all models that could be applied for that test pair. This design is particularly economical for sparse datasets as it provides for multiple uses of a single observation; a training pair can be used in training multiple models (e.g., pairs for which all features are present can be used for training all models). Given the large number of models to be trained (number of partitions times number of training folds), we did not explore optimizing random forest hyper-parameters; they were set to 400 trees and no maximum features limit. After completing all partition training, we generated a table of every partition’s estimated model accuracy, class imbalance weight, contributing features, and the number of contributing observations.

**Figure 1.**
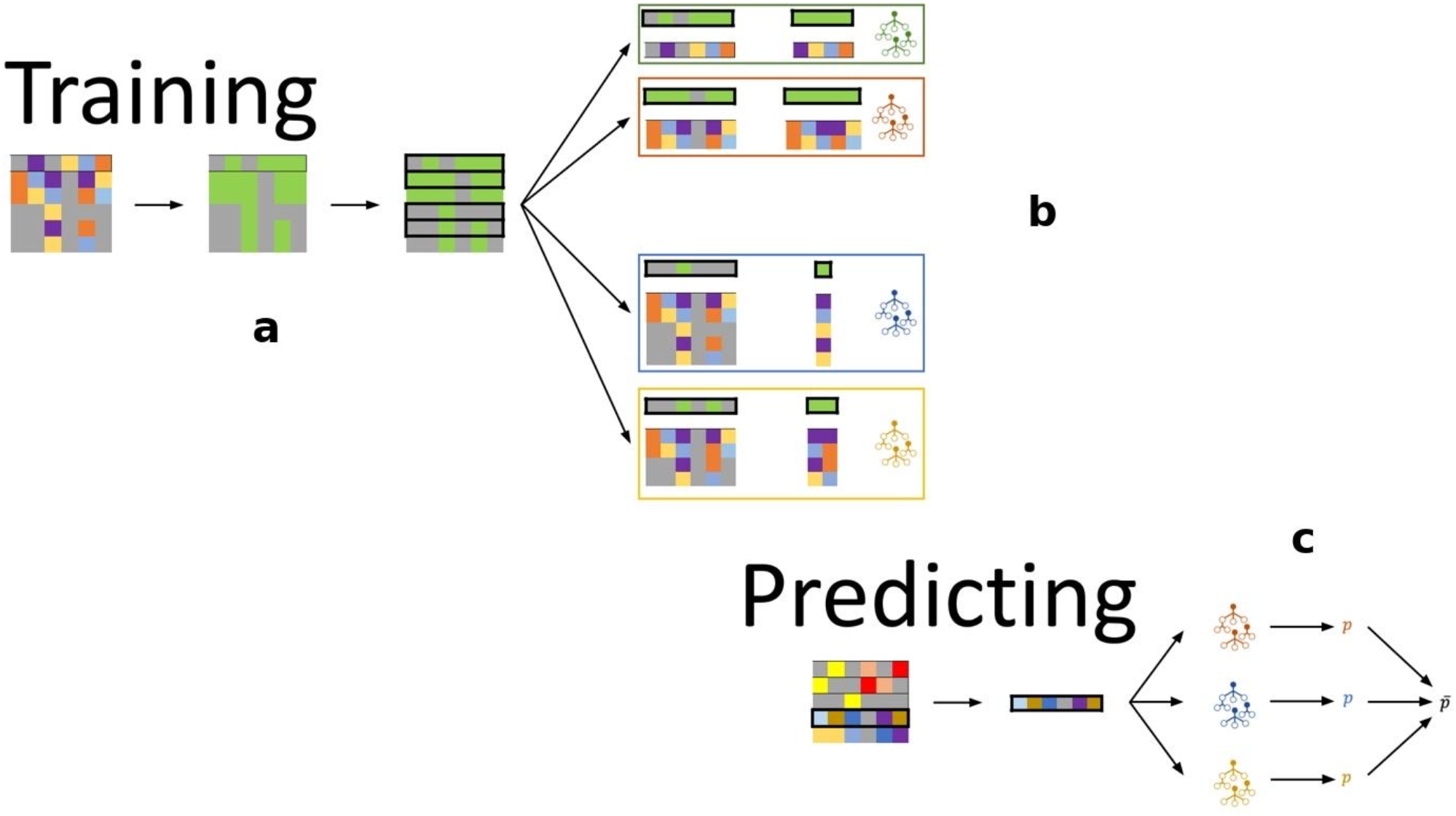
Overview of ELCFS approach. (a) identify all partitions of features for which there exists a real submatrix, e.g., having no missing data, (ii-b) learn a model for each partition, and (ii-c) calculate scores from the weighted average ensemble of each protein-pair.

### 2.2 Pair sets construction

Three sets of pairs were constructed for training models (see Table 1), including the features described in the next section where available. For these sets, a label of positive was assigned to a pair if both proteins are found in at least one complex in CORUM, and negative otherwise. The first pair set, referred to as D17, consisted of the union of the training and test sets of Drew et al., 2017 (Drew et al., 2017). This consists of 235,020 protein pairs and assigned labels, all of which have at least some features. It contains a mix of human and non-human proteins. The second (referred to as LM19) was the “expanded pairs” set described by Lugo-Martinez et al., 2019 (Lugo-Martinez et al., 2019). It consists of 1,038,969 pairs of only human proteins, all of which also had assigned labels. We constructed the third set, referred to as PS24, by starting with the human core set from the 01.07.2018 (3.0) version of the CORUM repository (Tsitsiridis et al., 2023b), which consisted of 3,375 proteins. All possible pairs of these were formed, producing 5,693,625 pairs. To these, 3,417,479 disjoint pairs from hu.MAP 1.0 and 403,242 disjoint pairs from hu.MAP 2.0 were added to give a set of 9,514,346 pairs (the inclusion of the hu.MAP pairs added 37,873,045 pairs that included 2,069 non-human proteins) (Drew et al., 2017, 2021). Of these, labels were available for 9,514,346 pairs, of which 6,601,161 pairs had features available.

**Table 1.** Properties of protein pair sets used in this study.

|  | hu.MAP1.0 | hu.MAP2.0 | LM19 | PS24 | STRING<br>v.12 |
| --- | --- | --- | --- | --- | --- |
| Pairs with data | 2,557,860 | 17,564,755 | 20,615,986 | 162,758,569 | 6,881,445 |
| Proteins with data | 13,276 | 15,554 | 6,883 | 19,340 | 19,232 |
| Possible pairs<br>(proteins with<br>data) | 88,119,450 | 120,955,681 | 23,684,403 | 187,008,130 | 184,925,296 |
| Pairs labeled | 5,446,857 | 3,435,064 | 1,408,681 | 9,514,346 | 6,184,140 |
| Proteins in labeled<br>pairs | 3,835 | 2,993 | 1,679 | 5,137 | 3,643 |
| Labeled pairs with<br>data | 235,020 | 2,299,721 | 1,038,169 | 6,601,161 | 773,026 |
| Fraction labeled<br>pairs predicted | 0.042 | 0.41 | 0.73 | 0.69 | 0.13 |
| Fraction proteins<br>with data predicted | 0.029 | 0.15 | 0.87 | 0.87 | 0.037 |
| Non-human<br>proteins | 1,800 | 583 | 0 | 2,069 | 0 |
| Human proteins<br>with data | 12,954 | 14,971 | 6,883 | 17,271 | 19,232 |
| Human possible<br>pairs | 83,896,581 | 112,057,935 | 23,684,403 | 149,135,085 | 184,925,296 |
| Non-human pairs | 1,619,100 | 169,653 | 0 | 2,139,346 | 0 |
| Human-non-human pairs | 20,656,800 | 8,728,093 | 0 | 35,733,699 | 0 |
| Complexes | 4,659 | 6,965 | 1,340/7,758 | 3,077 | 3,870 |
| Complexes'<br>proteins | 7,778 | 10,058 | 570/1,135 | 11,121 | 17,358 |
| Complexes' PPIs | 56,737 | 41,989 | 2,804/5,254 | 53,031 | 376,215 |
| Non-human<br>proteins in<br>complexes | 104 | 0 | 0 | 2 | 0 |

An additional pair set was constructed from the STRING version 12 release (Szklarczyk et al., 2023). It consisted of 19,232 proteins yielding 194,925,296 pairs. STRING is a database of known and suspected protein complexes derived from multiple sources, representing nearly 25 million proteins from over 5 thousand organisms. STRING provides multiple confidence metrics, including “combined score,” “text-mining” (from curated abstracts), “database,” “experimental” (directly measured experimental results), “co-expression” (from mRNA and DNA assays), “hscore” (from the degree of homology between potential interactors), “pscore” (from phyletic profiles), “fscore” (determined from fused proteins in specimens from different species), and “nscore” (calculated from inter-gene nucleotide counts). This set was used only for prediction.

### 2.3 Feature matrix assembly

The features used to predict PPI were pooled from various sources (see Table 2). They consisted of the 258 features used by Drew et al., 2017 (Drew et al., 2017) and 10 features derived from those of Lugo-Martinez et al., 2019 (Lugo-Martinez et al., 2019) (protein expression by tissue and by cell line were averaged to give one feature). To these, we added 1 feature measuring the correlation between transcript expression of a given pair using data from FANTOM5 (Lizio et al., 2015), the Genotype-Tissue Expression project (GTEx) (Consortium, 2020), and the study of Uhlen et al., 2015 (Uhlén et al., 2015). These are presumably normal tissue and, therefore, complement the NCI-60 data (Gholami et al., 2013) that comes from cancerous cell lines. This may help our results to have broader relevance across different cell states. An additional set of features was generated by applying a deep-learning-driven classifier (Ouyang et al., 2019) to the Human Protein Atlas images to assign a fraction of each protein found in 27 subcellular locations. We then calculated the correlation between those fractions for a given pair of proteins. This was done for each of the 5 cell lines. Finally, we added 5 features from the SubCellBarCode project in which cell extracts from 5 cell lines (A431, H322M, HCC827, MCF7, and U251) were fractionated into four subcellular compartments (secretory, nuclear, cytosolic, and mitochondrial) which were then subjected to CF-MS (Orre et al., 2019). Each feature represents the correlation between all the fractionation profiles for a protein pair in one cell line.

**Table 2.**
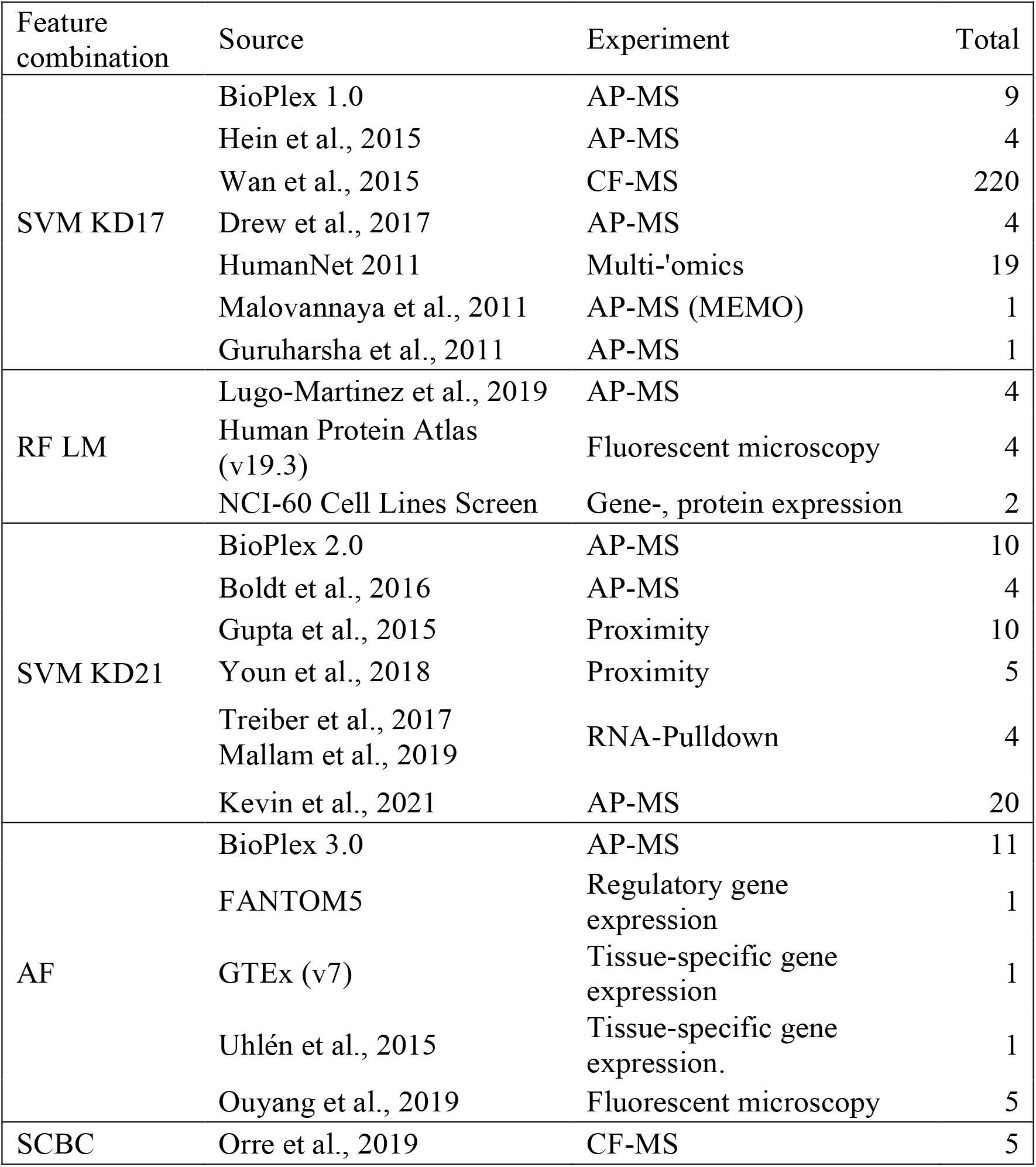
Feature types and sources used in different feature sets for predicting protein-protein interactions.

We also generated cell-type-specific feature values for three cell lines, H322M (lung), MCF7 (breast), and U251 (CNS). To do this, we replaced the correlation features for protein and RNA expression derived from NCI-60 with ratios of the values for a given pair of proteins for a specific cell line.

### 2.4 Interaction score prediction and evaluation

In all experiments, five-fold cross-validation of all labeled pairs for a given pair set was used for model learning, yielding a predicted interaction score for all labeled pairs. In addition, the trained model was used to predict interaction scores for all pairs for which a label was unavailable.

Due to heavy imbalance in the frequencies of positive and negative labeled pairs, we used the area under the precision-recall curve (AUC-PR) to estimate overall performance against labels generated from CORUM. However, some pairs considered non-interacting (negative label) because they are not present together in a CORUM complex may be present in complexes that have yet to be identified and added to CORUM. This would tend to artificially decrease measured accuracies. We, therefore, also performed precision-recall analysis using labels derived from STRING (Szklarczyk et al., 2023), which contains many more potential complexes. STRING contains metrics derived from various sources, as well as a combined score. For the combined score, we used the recommended threshold of 700 to assign labels. We did additional evaluations for the other metrics at thresholds for which the predictions elicited by the recommended “combined-score” were most similar.

### 2.5 Complex prediction and evaluation

Pairs were clustered into complexes using their predicted interaction scores and two-stage cluster discovery and refinement as described (Drew et al., 2021) (https://github.com/marcottelab/protein_complex_maps). Briefly, in the first stage, protein clusters are assembled one vertex at a time using a greedy algorithm (ClusterONE, https://github.com/ntamas/cl1) (Nepusz et al., 2012) until optimally cohesive, and in the second stage, they are subjected to Markov clustering (MCL) (https://github.com/micans/mcl) (Van Dongen, 2008) to extract the most probable embedded network. The cluster formation process required setting a number of hyperparameters. These included the initial PPI score threshold, the minimum density (the weighted sum of a cluster’s internal edges divided by its total possible edges), the max overlap allowed between clusters, whether to apply MCL and, if so, the inflation (cluster modularity strength) value. To choose appropriate ranges for these hyperparameters, we carried out preliminary clustering for a wide range and chose those values that maximized the accuracy of predicting ribosomal subunits (see Table S1). Complexes were generated for all 2,592 combinations of those values, and generated complexes were evaluated by comparison to CORUM (entire CORUM core set collection from the 01.07.2018 release) or STRING (all complexes in v12) complexes using the size weighted-F1 method described by Drew et al., 2017.

For the construction of consensus complex predictions, the eight hyperparameter combinations that optimized different criteria were chosen (see Table S2), and the unique complexes were combined. The criteria were (weighted F1 *or* weighted F1 x number of complexes) x (CORUM labels *or* STRING labels) x (without *or* with MCL).

Scores for individual predicted complexes within the merged set were assigned by two methods: taking the average or the minimum of the pair-wise PPI scores. Cell line-specific scores for each complex scores were assigned in the same manner using cell line-specific pair scores.

### 2.6 Data and software availability

A Reproducible Research Archive containing all software necessary to reproduce the results described here is available at https://github.com/GaryWilkins/elcfs_protein_complex_modeling. This includes scripts for downloading and preprocessing input data from various sources. The complete results of the processing and analysis are available in a public Google Drive folder.

## 3. RESULTS

As described in the Methods, we collected features derived from diverse sources useful for estimating the likelihood that protein pairs interact (see Table 2). There were extensive differences between sources regarding the protein pairs for which they provided values (illustrated in Figure 2A). For example, co-expression and co-localization measurements are available for almost all human protein pairs, whereas AP-MS features are only available for a relatively small fraction of pairs (Figure 2B). The resulting sparseness of the feature matrices creates a challenge for learning predictive models of pair interaction likelihood, and we, therefore, developed a novel variation of ensemble learning to deal with that sparsity. As described in the Methods, this approach (which we refer to as Ensemble Learning for Complete Feature Subsets, or ELCFS) learns separate models for sets of features that are shared by different sets of protein pairs. We applied this approach to increasingly larger sets of protein pairs and compared the results to those from prior studies.

**Figure 2.**
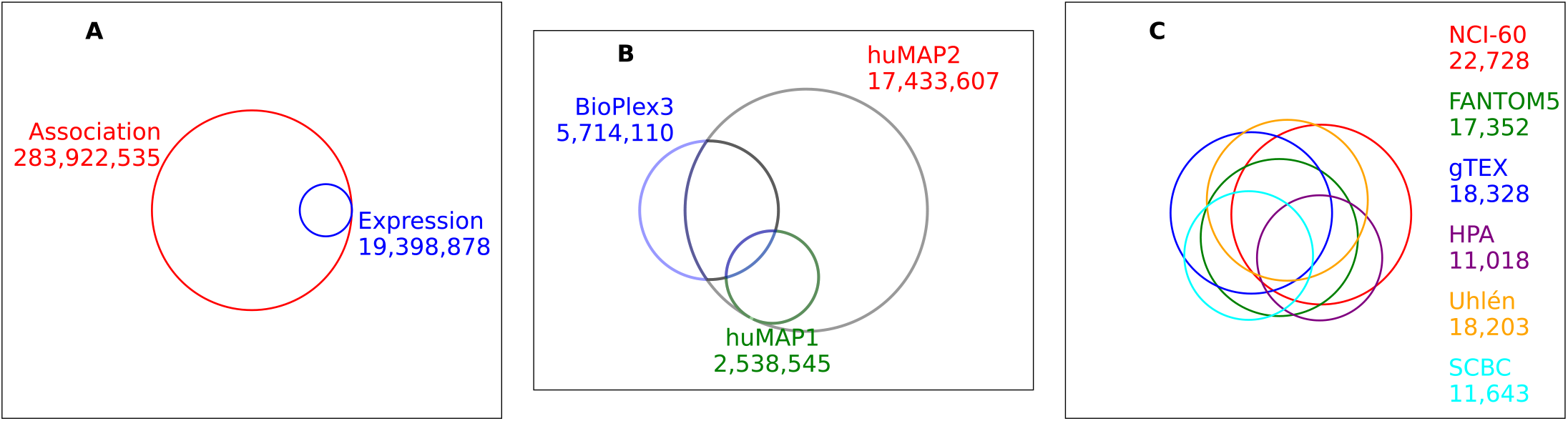
Venn diagrams of overlaps in feature sources and pairs sets used in this study.

## 3.1 Partition-based modeling improves performance with data from diverse experimental modalities

When analyzing a given set of protein pairs, it is important to distinguish between the proteins in that set, the pairs that those proteins could form, the subset of pairs for which experimental data is available, and the subset of pairs for which there is evidence on whether they form complexes. These subsets are expected to be overlapping but not identical.

We considered three sets of protein pairs in our study (see Table 1 and Methods). The first (referred to as D17) is the union of the protein pairs in the training and testing sets of the Drew et al., 2017 study (Drew et al., 2017). It consists of pairs for which both proteins were in the 08.10.2009 (2.0) version of the CORUM database (Ruepp et al., 2010) and for which AP-MS or fractionation features were available. The second (LM19) is the expanded pairs set defined by Lugo-Martinez et al. (Lugo-Martinez et al., 2019), which added NCI-60 and Human Protein Atlas (Uhlén et al., 2015) features in order to enable predictions for protein pairs that had not been observed in AP-MS or fractionation experiments. The third (PS24) consists of all pairs of human proteins for which gene expression or protein localization data were available, plus all pairs previously used by hu.MAP 1 and 2 (Drew et al., 2017, 2021): this set covers a larger number of human proteins than previous studies. The relationships between these sets are illustrated in Figure 2C.

For each pair set, it is important to note that the pairs that different sources provide data for may be biased toward those with positive (or negative) interactions. The features from AP-MS experiments, and the features prepared by Drew et al., 2017 (Drew et al., 2017) from the subcellular fractionation experiments of Wan et al (Wan et al., 2015), are both provided only for pairs that meet some threshold. This is why the number of pairs with data for the two hu.MAP studies (Table 1) is a small fraction of the possible pairs that could be formed between the proteins detected (this is also true for the pairs provided by STRING v12 (Szklarczyk et al., 2023), which are also thresholded). In contrast, the co-localization and co-expression features are available for almost all protein pairs, making the number of pairs with data in Table 1 much larger for the LM19 and PS24 sets.

We compared our modeling approach against previous studies for the different pair sets. The previous studies used separate training and test sets, while we chose to use cross-validation throughout our analyses. We divided the labeled pairs in a given set into 5 folds and used cross-validation to learn models and predict scores for pairs in each fold in turn. Pairs for which labels were unavailable were assigned the average score from the 5 models.

To generate comparable results, we applied the same strategy for the methods described previously (SVM for D17 and RF for LM19). Fig 2 shows precision-recall (Pr-R) curves and area-under-the-curve (AUC) values for the previous studies and various feature sets using our ELCFS approach.

For the D17 pair set (Figure 3a), the performances of previous approaches are similar, with the Drew21 performance being slightly lower. Using ELCFS with only the D17 features gave a similar AUC value, although incorrect high-scoring predictions lowered the early part of the curve. Note that as additional features were added, prediction performance improved strikingly. Using ELCFS with both the D17 and LM features yielded a boost of over 10%; and, including the SCBC features added another 10%. Improvement with SCBC features, which are similar to the CF-MS data from Wan et al (Wan et al., 2015), occurs presumably because of the additional information derived from multiple cell types and multiple subcellular fractions.

**Figure 3.**
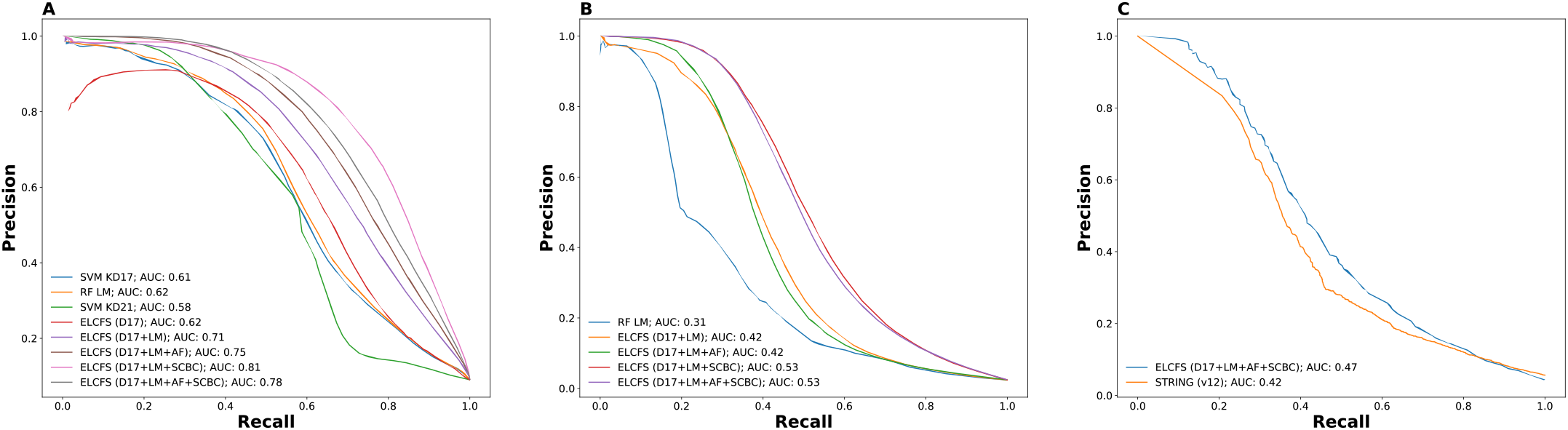
caption (Performance comparison for predictors of pairwise protein interactions. Precision-recall curves (and associated areas under the curves) are shown for different prediction methods and feature sets on (A) the D17 pair set described by Drew et al 2017 (Drew et al., 2017), (B) the LM19 pair set described by Lugo-Martinez et al (Lugo-Martinez et al., 2019), and (C) the PS24 pair set as defined in the Methods.

The results for LM19 are shown in Figure 3b. This set contains many pairs that were not in CORUM complexes and were therefore labeled as negative by default. However, at least some of these pairs may, in fact, be in previously undescribed complexes. The “false positives” may be the cause of the “apparent” AUC values (Figure 3b) that are lower than those for the D17 pair set. Nonetheless, the ELCFS method showed significant improvement over the LM19 approach for this set.

### 3.2 Comparison to STRING predictions

We also trained models using the labeled pairs in our PS24 set and assigned scores to all pairs. However, a comparison to previous approaches could not be done since they have not been applied to this larger set. Alternatively, we considered how our results compared to the more indirect interaction prediction approach used by the STRING database (Szklarczyk et al., 2023). STRING uses a variety of features, including literature co-mentions and database entries, to assign an interaction score to pairs of proteins. STRING v12 reports a pair as potentially interacting for those pairs scoring above 0.15. We therefore constructed the intersection of those STRING pairs and the PS24 set (which gave 6,870,437 pairs). Lastly, we constructed precision-recall curves (Figure 3c) for the subset of those pairs (approximately 19%, see Table 1) for which CORUM labels were available. As shown in Figure 3c, our approach yielded better results than STRING, at least as judged by CORUM labels.

We next compared predictions from hu.MAP 1 & 2 to those from our approach using the large set of pairs common to those studies and our PS24 set. Figure S1a shows precision-recall curves for the subset of those pairs that have CORUM labels. ELFCS provides more accurate predictions for this subset. We also evaluated the union of the pairs predicted by hu.MAP and by ELCFS. Since, for many of those pairs predictions are available from both approaches, we did the comparison two ways: giving preference to the hu.MAP score or to the ELCFS score. Figure S1b shows that ELCFS provides much more accurate predicted scores.

For all comparisons above, we used labels generated from CORUM as the ground truth. This, however, dramatically limits the number of predictions that can be evaluated. We, therefore, also used STRING as a source of ground truth. For this, we used three different thresholds on the STRING score to define positive or negative pair interactions. As shown in Figure 4a, the ELCFS predictions for the PS24 set match well to STRING scores, with AUC values up to 0.77 (note that this is consistent with the similar performance of ELCFS to STRING when used as a predictor and evaluated using CORUM labels in Figure 3c). When only the pairs that are contained in hu.MAP 1 and 2 are considered (Figure 4b), hu.MAP predictions perform dramatically worse than ELCFS.

**Figure 4.**
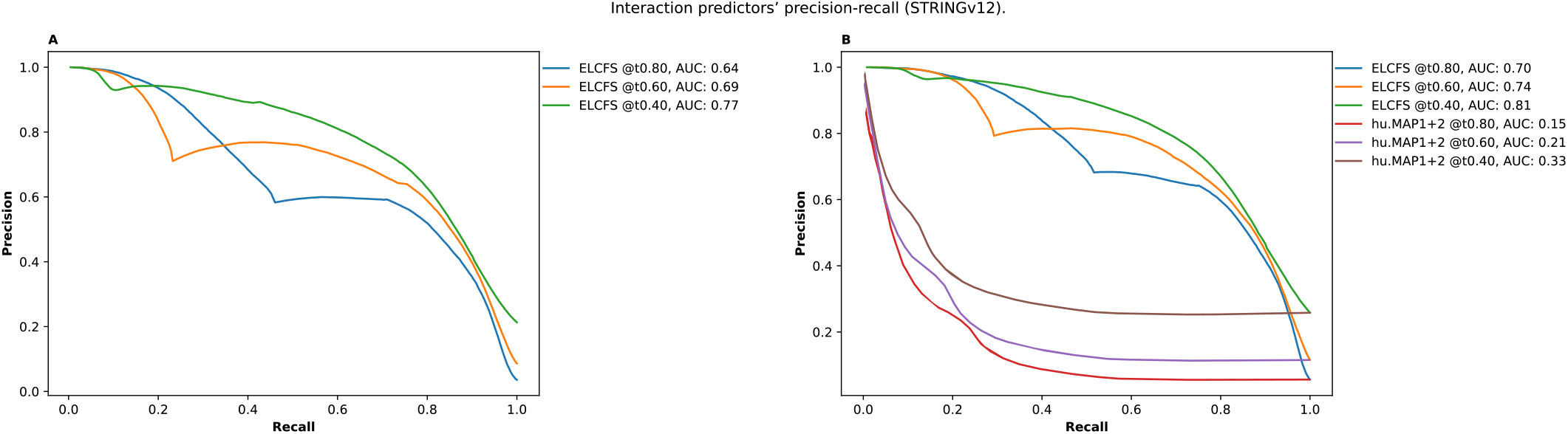
Interaction predictors’ precision-recall over STRING (v12). Performance comparison between predictors of pairwise protein interactions using labels derived from STRING (v12) confidence thresholds for ground truth. This figure shows the precision, recall and area under the curve (AUC) of our proposed method compared against previously published approaches over various pair sets, all within the intersection of STRING (v12) pairs: (A) PS24, (B) PS24 overlapping with hu.MAP 1+2, and (C) PS24 overlapping with hu.MAP 1+2 and Lugo-Martinez et al., 2019.

### 3.3 Construction of predicted complexes

Given the significantly improved performance of our new pair interaction prediction approach, we next used the pairwise scores to construct potential protein complexes using the methods previously described (Drew et al., 2021). Briefly, the approach consists of finding initial protein clusters (using ClusterONE) followed by optional refininement (using MCL).

As described in the Methods section, there are a number of hyperparameters that need to be chosen during this process. We therefore ran the complex assembly process for all combinations of a range of these hyperparameters. We evaluated the set of complexes from each hyperparameter combination (run) by calculating a weighted F1-score using either CORUM or STRING labels. Figure 5 shows results for each run for hu.MAP and ELCFS, divided into runs with or without using MCL after ClusterONE. There is significant variation across different hyperparameter combinations. As might be expected, there is a rough correlation between performance with CORUM or STRING labels. The best overall results, considering both labels, were achieved by ELCFS without MCL.

**Figure 5.**
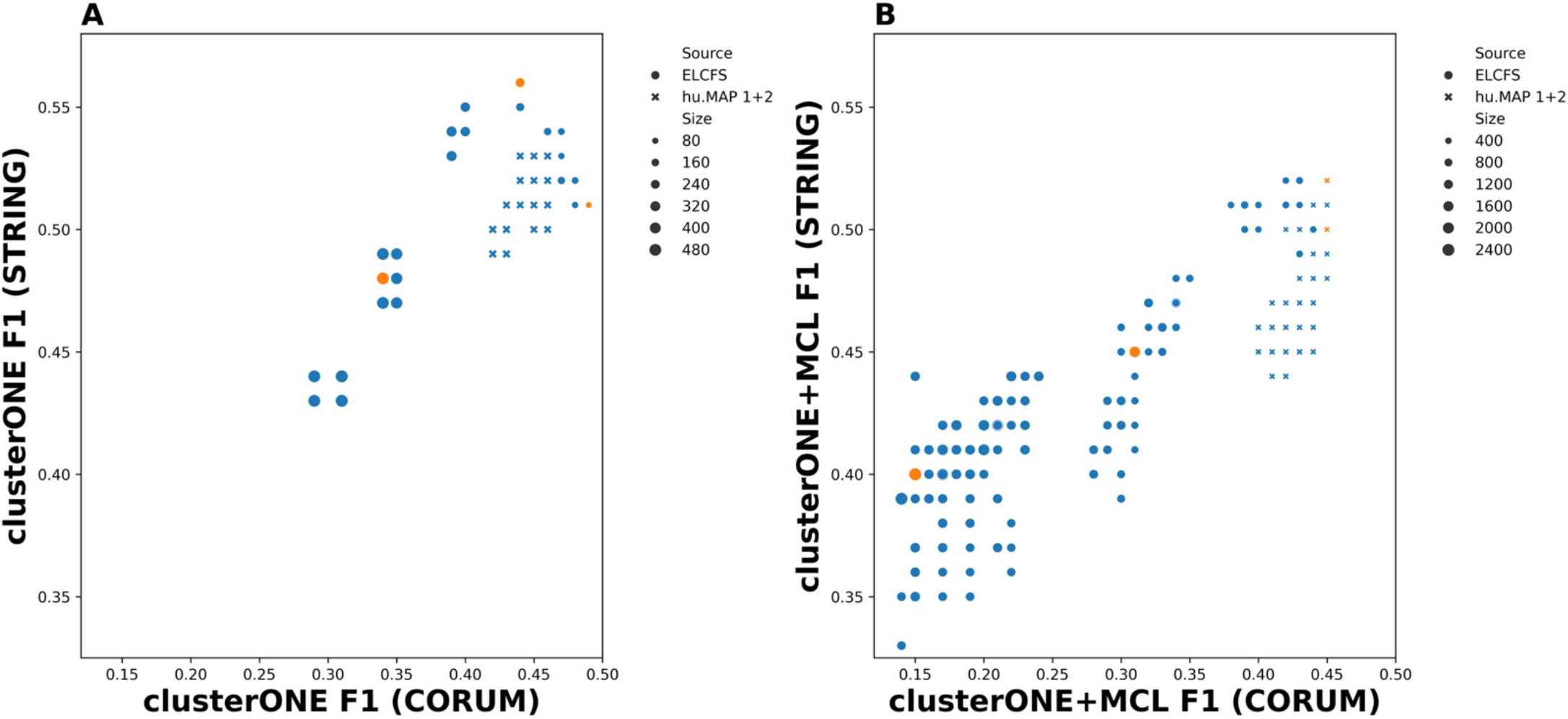
Comparisons of complex predictions relative to CORUM 01.07.2018 (3.1) or STRING v12. Symbols show complex assembly weighted F1-scores against both references for different assembly parameter sets either with or without MDL processing. The values for the eight assembly hyperparameter sets listed in Table S2 are marked in orange.

These weighted F1-scores reflect the overall performance of a hyperparameter set, and do not provide estimation of the likelihood of any particular complex. To estimate this, we calculate both the minimum interaction score for any pair (a very conservative measure that requires evidence of interaction for all pairs) and the mean interaction score over all pairs.

The different hyperparameter combinations emphasize different aspects of complex formation, primarily trading off between size and stringency (confidence). We therefore chose to construct a set of consensus complexes (which we term CS24) by merging the results from eight hyperparameter combinations that optimize different criteria (see Table S2 and Methods). This resulted in a total of 1700 predicted complexes. A detailed analysis of this large set is beyond our scope here. We therefore discuss selected sets of illustrative complexes that either closely match CORUM complexes or are novel.

The addition of features that try to provide more indirect evidence of protein-protein interaction may be expected to slightly expand the membership of currently known complexes. For example, proteins that are actually part of a complex may not be included in CORUM (e.g., because their peptide composition makes them harder to detect in affinity-purification mass spectrometry experiments) but may be included in our predictions due to supporting co-expression or co-localization data that raises the probability of interaction. Conversely, membership of known complexes that are regulated (e.g., by subtracting specific members to reduce activity that might reduce co-expression) may be expected to shrink if it results in lower estimated interaction probability for certain pairs. Both phenomena occur frequently in our predicted complexes, and only 4 predicted complexes matched exactly with a CORUM complex (Table S3). Table 3a shows five predicted complexes that have 1-3 fewer proteins than a CORUM complex but no additional proteins. For example, one of these (complex 1971) is the exosome complex minus EXOSC6 and CD-1. This suggests that different subsets of the complex may be present. Table 3b shows the opposite case, three predicted complexes containing 12 proteins that exactly match a known complex (the multisynthetase complex) but each having one additional protein.

**Table 3.**
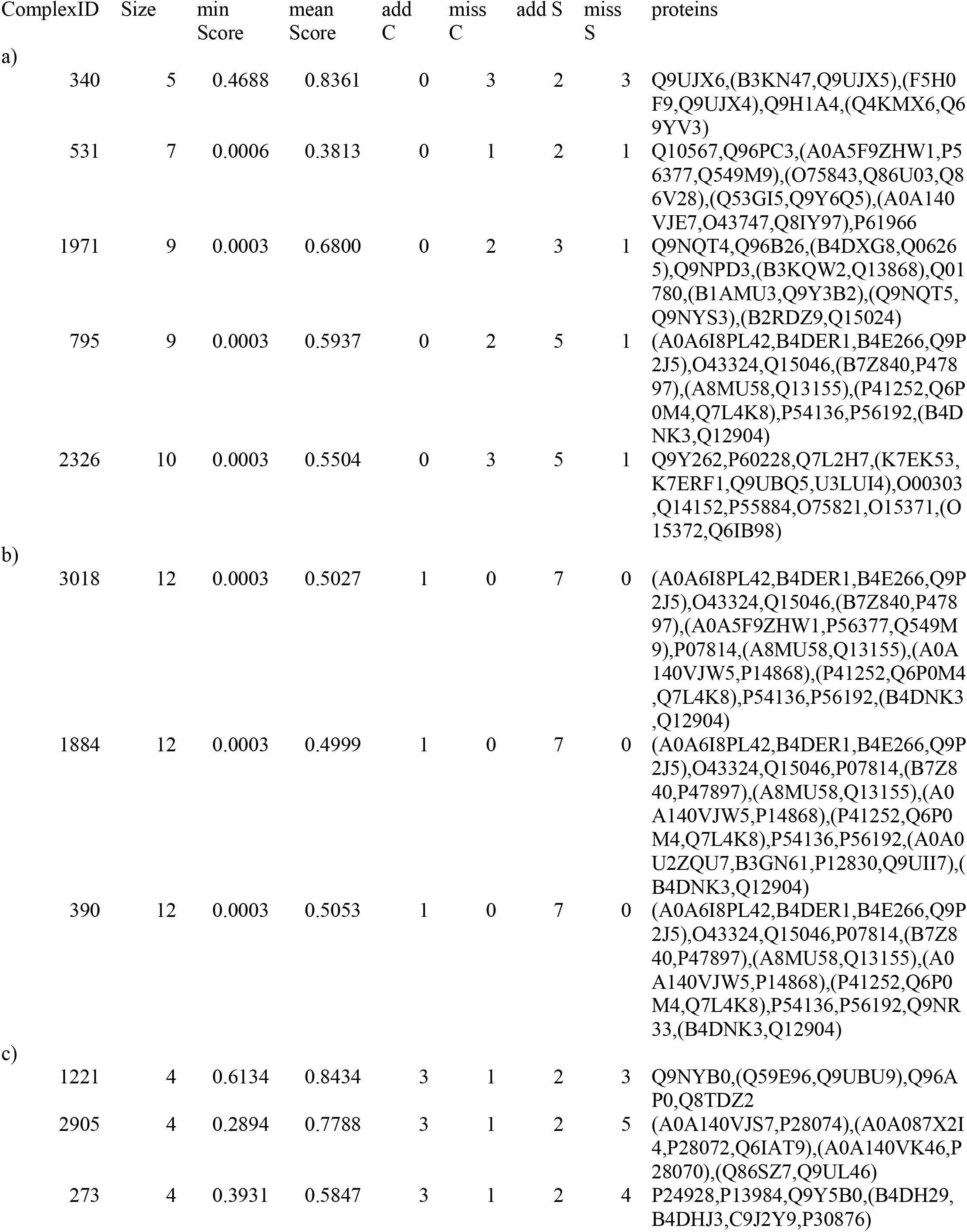
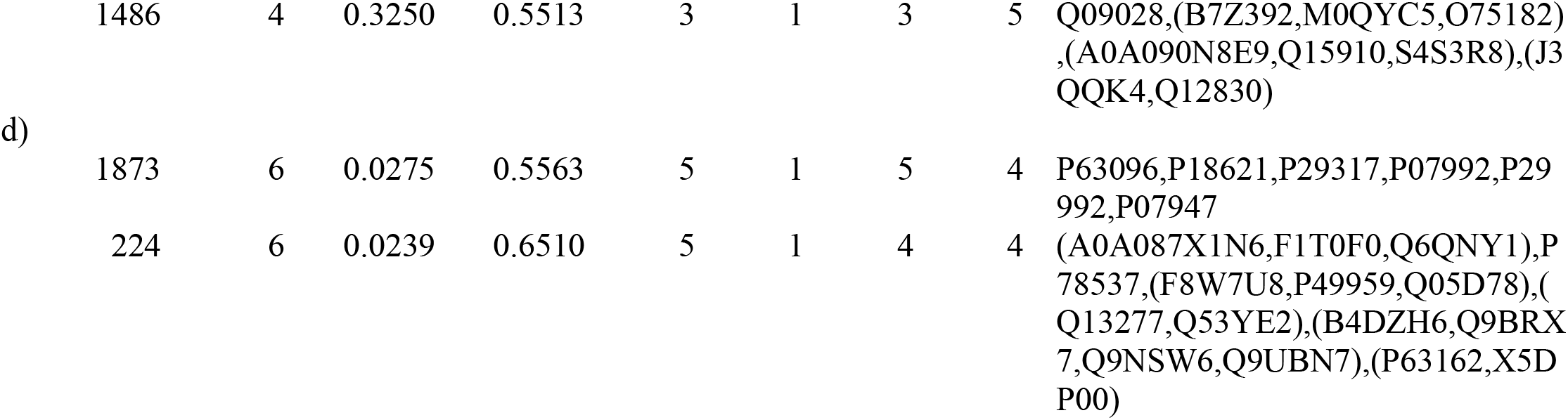
Selected predicted complexes. addC and missC refer to number of additional and missing proteins compared with the most similar CORUM complex, addS and missS refer to differences with the most similar STRING complex.

Of course, the broader coverage of the proteome that our features provide is also expected to yield novel predicted complexes. For example, Table 3c shows four novel predicted complexes of length 4 that had very high estimated likelihoods using both the minimum and mean pairwise interaction score. These involve proteins with diverse known functions and are considered examples of what Drew et al., 2017, 2021 referred to as “promiscuous proteins.” As another example, Table 3d shows two novel complexes of length 6 that scored high in mean pairwise score but low in the minimum pairwise score. This occurs when interaction scores are high for some but not all of the pairs, perhaps because of differences in the coverage of the proteins between different data sources.

### 3.4 Cell line-specific PPI models

Changes in the behavior of different cell types presumably come, at least in part, by up- or down-regulation of the formation of different protein complexes. Since the detailed AP-MS experiments in the BioPlex project have not been performed in parallel for different cell types or cell lines, we considered whether we could at least make initial predictions of differences in protein interactions between cell types using our approach. To do this, we constructed two cell-line specific features to reflect the relative expression of pairs of proteins. As described in the Methods, we calculated the ratios for protein or RNA expression in the NCI-60 dataset, and normalized them to match the ranges of the correlation features used previously. This was done for three cell lines derived from different tissues, H322M (lung), MCF7 (breast), and U251 (CNS). We used our trained (non-cell line specific) models to make predictions for each pair for each line after replacing the two NCI-60 co-expression features with the cell line-specific versions.

Figure S3 compares the predicted PPI scores between pairs of those cell lines. As expected, most protein pairs have low scores, and those scores vary quite a bit between the cell lines. In contrast, the small fraction of very high scoring pairs (greater than 0.9) have similar scores for the cell lines, especially for H322 and MCF7. However, many pairs with medium to high scores (0.5 and higher) vary quite dramatically between cell lines. To assess this further, Table S4 summarizes what fraction of predicted PPIs scoring above a threshold of 0.28 were exclusive to each cell line or shared between and across the others. Though nearly 75% of each cell line’s above threshold predictions were shared with the other cell lines, we indeed found that there were predictions exclusive to each of the cell lines (with almost twice as many for U-251).

### 3.5 Cell line-specific complex predictions

The cell line-specific PPI scores allowed us to consider potential differences between cell lines in the CS24 consensus complexes described above. For this, two scores were generated for each complex for each cell line, one using the mean of the PPI scores and one using the minimum.

Note that calculating cell line-specific scores for the already identified complexes is different from generating complexes from scratch using cell line-specific PPI scores. Since building complexes considers only pairs scoring above an assembly threshold, the latter approach could potentially magnify slight differences in PPI scores between cell lines and result in different sets of complexes for each cell line. (For example, if a pair score for one cell line is just above the assembly threshold while just below for another, it would be potentially included in a complex for the first but not the second.)

Figure 6 shows that the variation in predicted complex mean PPI scores among the cell lines is much less than the dramatic variation in individual pair interaction scores among the cell lines (Figure S3). The Pearson correlation coefficient between the mean scores for pairs of cell lines ranges from 0.9935 to 0.9957; there is slightly more variation in complex minimum PPI scores, with the correlation ranging from 0.9882 to 0.9914.

**Figure 6.**
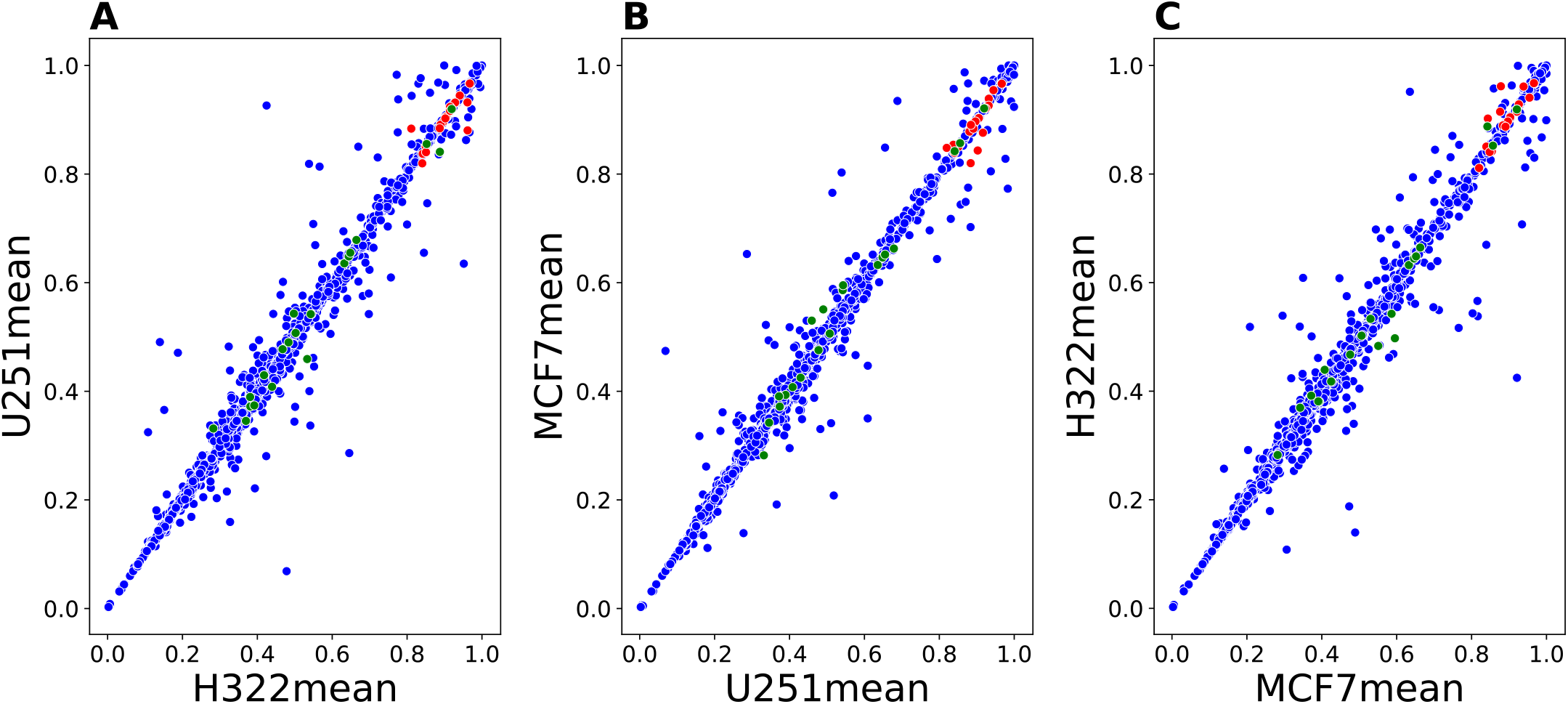
Comparison of complex scores for different cell lines. The scores for the 20 highest scoring complexes from non-cell specific predictions are shown in red and the scores for the 20 complexes most closely matching CORUM complexes are shown in green.

Similar to Table S4, Table S5 summarizes the fractions of predicted complexes that were exclusive, shared, or in common across the three lines. It uses a fixed, somewhat conservative, threshold to decide whether a complex is likely to be present in a given line. The number of predicted differences are quite small.

An alternative approach is to identify complexes with the largest differences in predicted scores between any pair of cell lines, which does not require a threshold. Table 4a shows two example predicted complexes that match CORUM complexes but have a difference in predicted score of close to 0.2 for one cell line compared to the other two. The first of these complexes has three out of four proteins in the TIN2 complex, which is a subset of both the telomere-associated protein complex and the RAP1 complex. Some of these proteins are found in four other CORUM complexes with different additional proteins. The second matches with five different variations on the APC/C (anaphase-promoting) complex. Both TIN2 and APC/C being found in multiple versions suggest that they may be regulated by addition or subtraction of subunits; the differences in the predicted scores for these complexes may reflect differences in the regulation of those complexes between the cell lines.

**Table 4.** Example protein complexes with differences in predicted score between different cell lines. Columns “min Score” and “mean Score” refer to the scores predicted using non-cell line specific features. Column “add C” and “mis C” refer to the number of proteins that a given complex has that are not included in the matching CORUM complex and the number of proteins that are present in that CORUM complex and not present in the predicted complex. “add S” and “mis S” refer to the analogous differences with the matching STRING complex. The columns under cell line names contains predicted cell-line specific scores. The last column contains the UniProt identifiers of the proteins in a given complex (with the identifiers in parentheses being alternatives from the same gene). a) Predicted complexes matching CORUM complexes. b) Predicted complexes whose proteins are not found together in a CORUM complex.

| Complex ID | Size | min Score | mean Score | add C | miss C | add S | miss S | H322 mean Score | MCF7 mean Score | U251 mean Score | proteins |
| --- | --- | --- | --- | --- | --- | --- | --- | --- | --- | --- | --- |
| a) |  |  |  |  |  |  |  |  |  |  |  |
| 2472 | 3 | 0.285 | 0.508 | 0 | 1 | 0 | 2 | 0.221 | 0.415 | 0.221 | (A8MTK3,Q5MJ33,Q9NUX5),Q96AP0,Q15554 |
| 340 | 5 | 0.469 | 0.836 | 0 | 3 | 2 | 3 | 0.576 | 0.410 | 0.421 | Q9UJX6,(B3KN47,Q9UJX5),(F5H0F9,Q9UJX4),Q9H1A4,(Q4KMX6,Q69YV3) |
| b) |  |  |  |  |  |  |  |  |  |  |  |
| 1629 | 3 | 0.363 | 0.658 | 2 | 3 | 1 | 3 | 0.663 | 0.215 | 0.210 | (I3L3E8,Q9P086),(O43513,Q6IAZ5),Q6P2C8 |
| 1438 | 3 | 0.191 | 0.379 | 2 | 1 | 2 | 4 | 0.091 | 0.085 | 0.398 | P07949,(M0R1I2,Q03405),P67809 |
| 736 | 3 | 0.401 | 0.542 | 2 | 1 | 0 | 11 | 0.570 | 0.320 | 0.313 | (P29374,Q05CG0),(Q13547,Q6IT96),(G5E9I4,Q9HCU9) |
| 2662 | 3 | 0.191 | 0.529 | 2 | 1 | 2 | 5 | 0.058 | 0.258 | 0.258 | O14745,(B2RCT6,P15170,Q7KZX8),Q13490 |
| 2901 | 3 | 0.743 | 0.753 | 2 | 1 | 2 | 4 | 0.742 | 0.559 | 0.728 | Q9H211,P38936,(P61024,Q5T178) |

Table 4b shows five examples of predicted complexes that do not match CORUM complexes and show differences in predicted score for different cell lines of as much as 0.45. The first contains three proteins that are found in various transcriptional regulator complexes in CORUM but are not found together. All three are, however, found together in a HuMAP2 predicted 32-protein complex (HuMAP2_01210) that consists of RNA Polymerase II mediators. Again, our predictions suggest differences in the regulation of the composition of mediator complexes between the cell lines.

## 4. CONCLUSIONS

Our method integrates data from orthogonal experimental sources to better predict protein-protein interactions. It successfully addresses two challenges. The first is that considerable differences in the coverage of proteins in different sources leads to widely different numbers of features available for each protein pair, making training of predictive models difficult. This has been addressed in the SVM-based models of Drew et al. (2017, 2021) by replacing missing features with zeros, but this introduces bias. The problem becomes worse when expanding the protein pairs to include those with only a few features, and feature inference cannot help because there is so little information to based it on. We therefore implemented a data partitioning scheme to better handle the sparse nature of the relevant available features. This trains many learners that each are based on a subset of protein pairs for which a subset of features are fully available. This allowed us to significantly improve the accuracy of our predictions over prior work without acquiring more data.

The second challenge is to address differences in protein interactions and complexes between different cell lines, types and tissues. Collecting AP-MS data for each of these is infeasible. Since our models include features that measure coexpression and colocalization, which are readily available or may be easily acquired, we propose that these be used to make predictions of the probability of a given protein interaction for a given cell type or tissue. We demonstrate this for three cultured cell lines but by substituting cell line-specific values for RNA and protein co-expression. We found that even small changes in feature values from cell type to cell type can produce large differences in predicted occurrence of specific protein interactions, but that in this test case produced modest differences in predicted complexes.

It is important to note that all of the work on predicting protein-protein interactions is subject to the effects of bias in the proteins for which experiments have been done. For example, the selection of proteins for AP-MS experiments may be affected by the proteins that have been previously characterized and for which suitable antibodies are available.

We suggest that our approach for making can cell type-specific predictions can conceivably be used to make initial predictions for a range of cell types and that active machine learning can then be used to selectively choose additional experiments (e.g., AP-MS) to iteratively improve the predictive models.

## 5. ACKNOWLEDGMENTS

This work was supported in part by NIH grants GM103712.

