## Supplementary Information for "Improved protein interaction models predict differences in complexes between human cell lines"

Gary R. Wilkins, Jr., Jose Lugo-Martinez* and Robert F. Murphy*
Ray and Stephanie Lane Computational Biology Department, School of Computer Science, Carnegie Mellon University

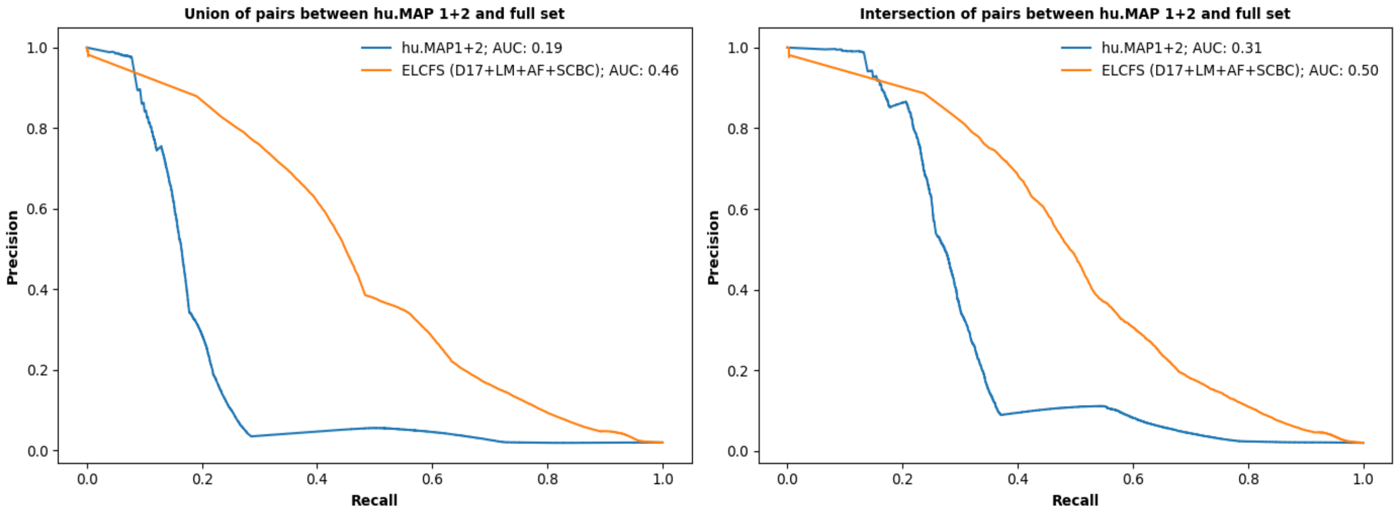


Figure S1. Performance comparison of merged results for different predictors. Predictions from each predictor were combined and results for one or the other model were used when there was overlap. Results are shown for A) the union of pairs in PS24 and hu.MAP 1+2, and B) the intersection of PS24 and hu.MAP 1+2. The legend shows which predictor was given preference for overlapping pairs.

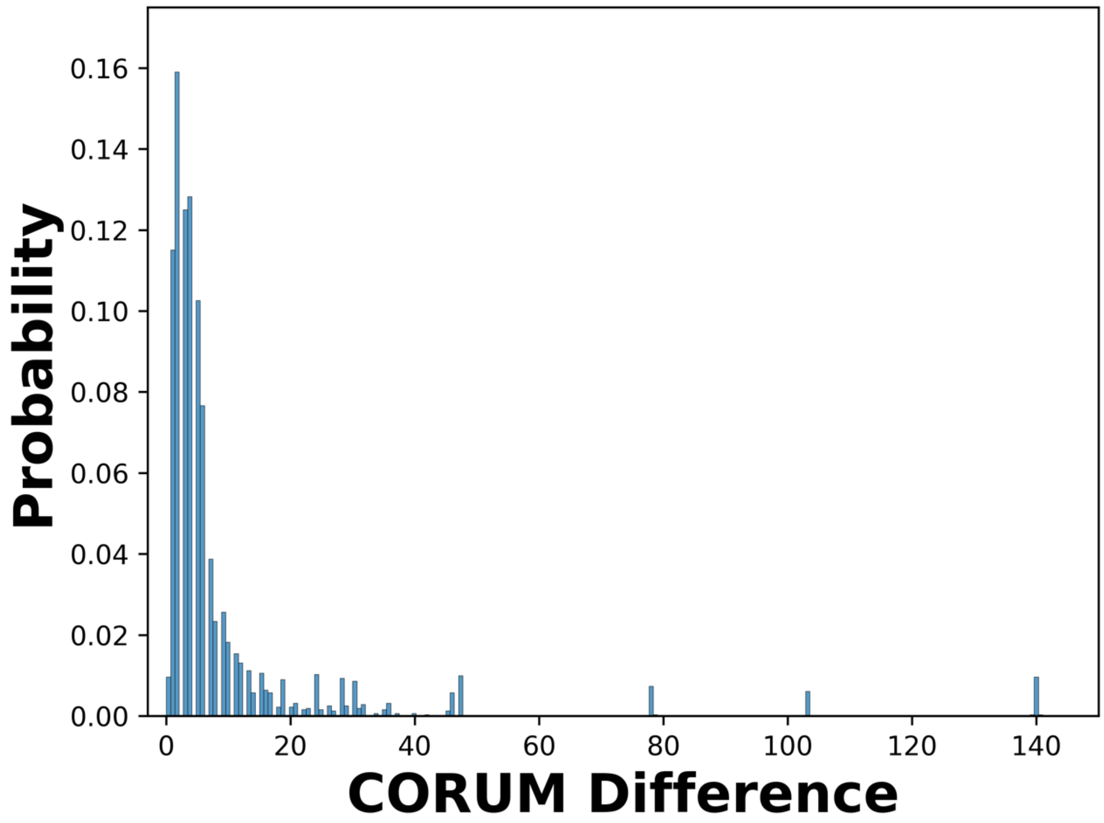


Figure S2 Differences in number of proteins between each CS24 complex and the closest matching CORUM complex. Note that most differences are only a few proteins.

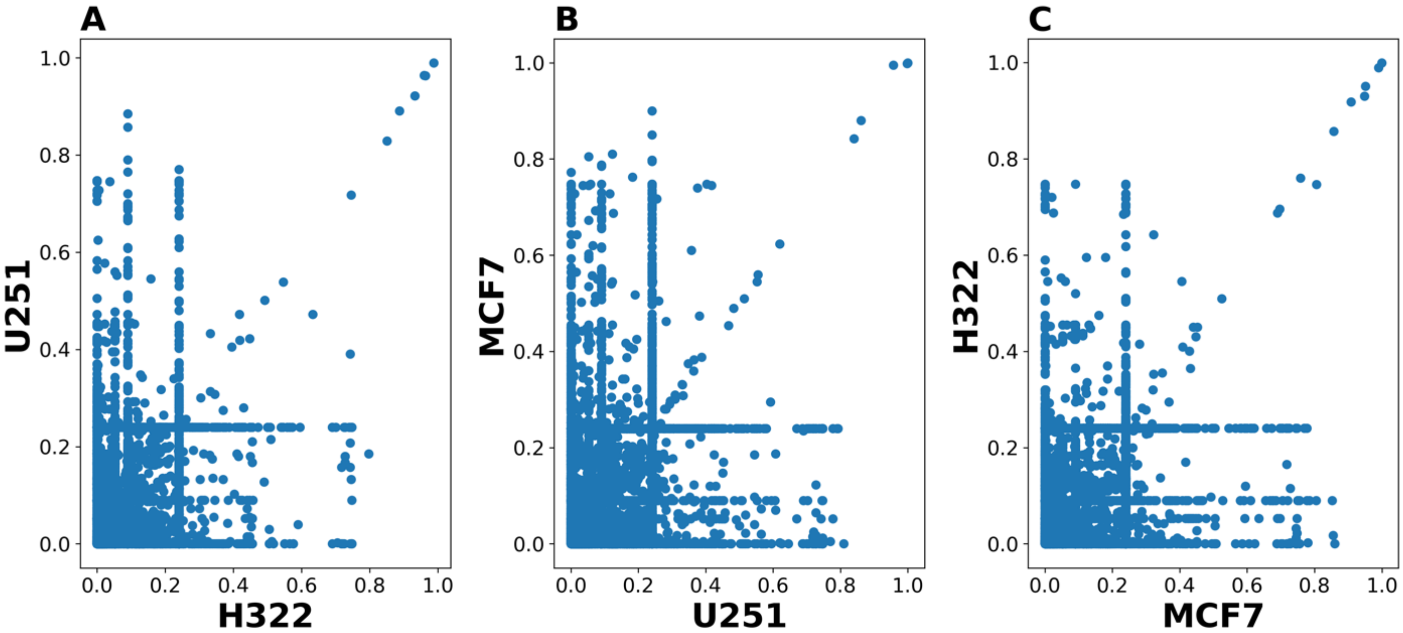


Figure S3 Comparison of predicted protein-protein interaction scores between pairs of cell lines. Given the large number of pairs, the scatterplots show only a randomly chosen subset of protein pairs. Note that low-scoring pairs have greater variability.

| | | | | | | |
|---|---|---|---|---|---|---|
| **threshold** | None | 0.267 | 0.280 | 0.293 | 0.307 | 0.320 |
| **density** | 0.025 | 0.075 | 0.100 | 0.200 | 0.300 | 0.400 |
| **max overlap** | 0.200 | 0.300 | 0.400 | 0.500 | 0.600 | 0.700 |
| **inflation** | 2 | 4 | 6 | 7 | 8 | 9 |

Table S1. Complex assembly parameter values used in this study. Complexes were built using all combinations of these values.

| Predictor | Thresh(s) | Method | Overlap | Density | Inflation (MCL) | Ref. | Metric | Score | Total |
|---|---|---|---|---|---|---|---|---|---|
| ELCFS | 0.28 | clusterONE | 0.2 | 0.02 | | C | F1*Total | 0.49 | 58 |
| ELCFS | 0.28 | clusterONE w/ MCL | 0.2 | 0.02 | 2 | C | F1 | 0.44 | 436 |
| ELCFS | 0.28, 0.29, 0.31, 0.32, None | clusterONE | 0.7 | 0.3 | | C | F1*Total | 163.54 | 481 |
| ELCFS | 0.28 | clusterONE w/ MCL | 0.3 | 0.02 | 4 | C | F1*Total | 566.06 | 1826 |
| ELCFS | 0.28, 0.29, 0.31, 0.32, None | clusterONE | 0.7 | 0.1 | | S | F1 | 0.56 | 239 |
| ELCFS | 0.28 | clusterONE w/ MCL | 0.5 | 0.05 | 2 | S | F1 | 0.52 | 541 |
| ELCFS | 0.28, 0.29, 0.31, 0.32, None | clusterONE | 0.7 | 0.3 | | S | F1*Total | 230.88 | 481 |
| ELCFS | 0.28 | clusterONE w/ MCL | 0.3 | 0.02 | 9 | S | F1*Total | 986.8 | 2467 |

Table S2. Combinations of assembly parameters providing best performance by various measures. The ‘Ref’ column shows the reference complexes were compared to, either CORUM (‘C’) or STRING (‘S’). The ‘Metric’ column shows how the parameter combination in that row was chosen as the best: ‘F1’ refers to the weighted F1-score of the predicted complexes against the reference which was optionally multiplied by the total number of complexes (‘Total’).

| ComplexID | Size | min Score | mean Score | add S | miss S | proteins |
|---|---|---|---|---|---|---|
| 2250 | 3 | 0.246 | 0.557 | 0 | 8 | Q13952,P23511,(F8VSL3,P25208) |
| 953 | 3 | 0.181 | 0.508 | 0 | 7 | Q9UPZ3,(G5E9V4,Q969F9),Q86YV9 |
| 1836 | 3 | 0.054 | 0.299 | 0 | 2 | (A0A1W1GSK9,A0A804HKF9,B4DXD8,Q92889),P07992,P23025 |
| 769 | 4 | 0.002 | 0.634 | 1 | 2 | Q9Y248,(A0A0S2Z5L0,A0A0S2Z5L4,A0A0S2Z5P2,Q9BRX5),Q9BRT9,(A0A8Q3WLK7,Q14691) |

Table S3. Predicted complexes exactly matching CORUM complexes. “add S” refers to the number of additional proteins in the predicted complex compared to the closest matching STRING complex, and “miss S” refers to proteins missing in the predicted complex compared to that closest match.

| First cell line | Total pairs predicted above threshold | Common to all cell lines | MCF7 | H322 | U251 |
|---|---|---|---|---|---|
| MCF7 | 2,189,452 | 1,663,513 | 325,930 | 86,004 | 114,005 |
| H322 | 2,281,754 | 1,663,513 | 86,004 | 363,950 | 168,287 |
| U251 | 2,763,480 | 1,663,513 | 114,005 | 168,287 | 817,675 |

Table S4 Distribution of protein pairs with high predicted interaction scores among cell lines. The second column gives the number of pairs whose mean score for the cell line in that row were higher than a threshold (0.28). These were separated into those common to all cell lines, and those in common with each other cell line, and those unique to that cell line (shown in the column matching the row).

| First cell line | Total complexes predicted | Common to all cell lines | MCF7 | H322 | U251 |
|---|---|---|---|---|---|
| MCF7 | 436 | 414 | 9 | 6 | 7 |
| H322 | 435 | 414 | 6 | 9 | 6 |
| U251 | 432 | 414 | 7 | 6 | 5 |

Table S5. Distribution of high confidence complexes among cell lines. The second column gives the number of complexes whose mean score for the cell line in that row were higher than a threshold (0.5527). These were separated into those common to all cell lines, and those in common with each other cell line, and those unique to that cell line (shown in the column matching the row).
